# Remus: System for Remote Deep Brain Interventions

**DOI:** 10.1101/2021.05.05.442844

**Authors:** Taylor D. Webb, Matthew G. Wilson, Henrik Odéen, Jan Kubanek

## Abstract

**Motivation:** Transcranial focused ultrasound brings personalized medicine to the human brain. Ultrasound can modulate neural activity or release drugs in specific neural circuits but this personalized approach requires a system that delivers ultrasound into specified targets flexibly and on command.

**Summary:** We developed a remote ultrasound system (Remus) that programmatically targets deep brain regions with high spatiotemporal precision and in a multi-focal manner. We validated these functions by safely modulating two deep brain nuclei—the left and right lateral geniculate nucleus–in a task-performing non-human primate. This flexible system will enable researchers and clinicians to diagnose and treat specific deep brain circuits in a noninvasive yet targeted manner, thus embodying the promise of personalized treatments of brain disorders. Animated graphical abstract: onetarget.us/download/rem.

## Introduction

One in three patients across brain disorders is treatment-resistant^1–8^. Neuromodulation provides these patients with new treatment options, but invasive, surgical approaches are risky and inflexible, and current noninvasive approaches do not have the necessary spatial resolution. Transcranial focused ultrasound provides a new set of regimens that enable the manipulation of brain activity noninvasively and at high spatiotemporal precision. Depending on stimulus duration, ultrasound modulates neural activity^9–14^ or induces changes in functional connectivity^15–17^. In addition, when combined with nanoparticles or microbubbles, ultrasound can be used to deliver drugs, genes, or stem cells selectively into the specified target/s. This can be achieved by releasing drugs from nanoparticle carriers,^18–20^ or using microbubbles to temporarily disrupt the blood brain barrier^21^ and so deliver large agents that would not pass otherwise. Thus, this form of energy provides novel therapeutic options for the millions of patients who are currently not adequately treated.

Nonetheless, these applications are limited by technological challenges. A key requirement for the success of the ultrasound-based therapies is the ability to target a specified site flexibly and reproducibly. This flexibility is critical given that the neural sources of many mental and neurological disorders are poorly understood and vary from individual to individual^22–25^. Systems that could provide this functionality— phased arrays—are primarily designed for ablative treatments^26^ with only a few phased array systems available for other transcranial applications^27,28^.

We developed Remus, a system that manipulates deep brain circuits remotely and programmatically, thus enabling personalized circuit manipulations in the clinical and research settings. The platform enables an operator to specify one or more targets, together with the desired timing and ultrasonic waveform, in software. The design is MR-compatible, which enables clinicians and researchers to confirm precise targeting. The system’s imaging functionality ensures reproducible positioning of the device and evaluates the quality of the ultrasound coupling to a subject’s head. We validated these capabilities in a nonhuman primate (NHP). Remus can be used to validate existing and devise new ultrasonic protocols in large animals, NHPs, and humans. Ultimately, the system will be used to provide precision treatments to patients who are currently out of options.

## Results

Remus is a practical and affordable platform that delivers focused ultrasound into specified deep brain targets remotely, with the skull and skin intact (Figure 1, top). The targets of the ultrasound are specified in software. The software can flexibly target individual sites in sequence (Figure 1, bottom) or simultaneously. We validated these features in a NHP. Specifically, we used the system to programmatically deliver focused ultrasound of specific parameters into the left and right LGN while quantifying the effects of the ensuing neuromodulation on behavior (Figure 1, top). The two LGNs were targeted electronically, without mechanical movement of the transducer.

**Figure 1.**
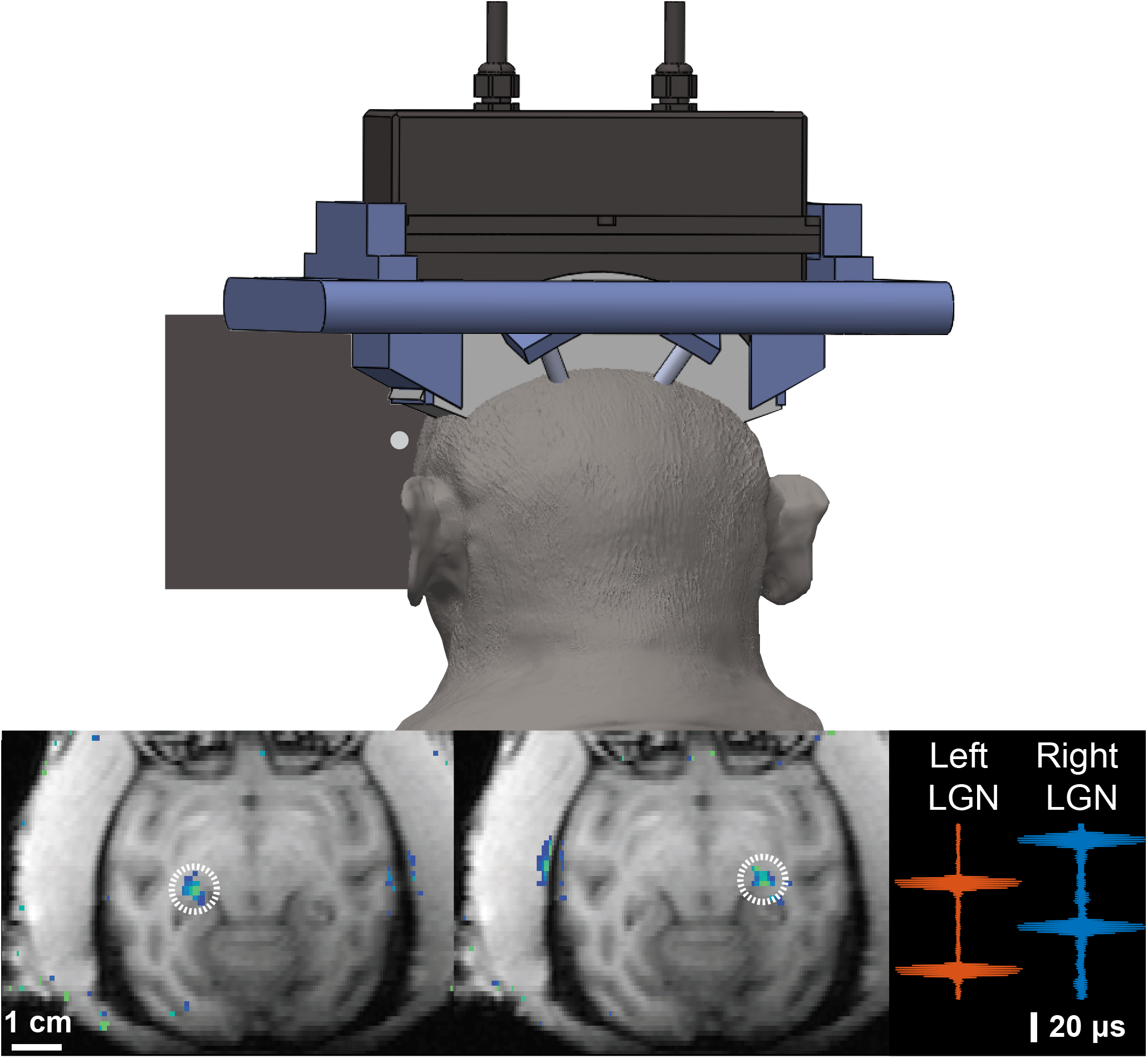
Remus: Remote ultrasound system for multifocal deep brain interventions. **Top:** The system is attached to the head of a NHP to provide reproducible targeting of deep brain circuits while the subject engages in behavioral tasks. **Bottom Left:** Ultrasound targets are specified programmatically. This figure shows selective targeting of the left and right lateral geniculate nucleus (LGN). Image overlays were acquired using MRI thermometry (see Methods). **Bottom Right:** The system can be used to target multiple sites in rapid succession (minimum pulse separation of approximately 30 *μs*) or simultaneously. The figure shows acoustic traces simultaneously recorded by two hydrophones placed at the approximate location of the left and right LGN.

We found that brief, 300 ms pulses of ultrasound directed into the left and right LGN transiently modulated choice behavior in the NHP. The subject was asked to look at a left or a right target, whichever appeared first. We quantified effects of the ultrasonic neuromodulation in the controlled condition in which both targets appeared at the same time. We assessed the proportion of choices of each target when the left LGN was stimulated, when the right LGN was stimulated, and quantified the proportion of choices of the contralateral target for these conditions. The resulting proportion of contralateral choices, averaged across individual sessions, is shown in (Figure 2, top).

**Figure 2.**
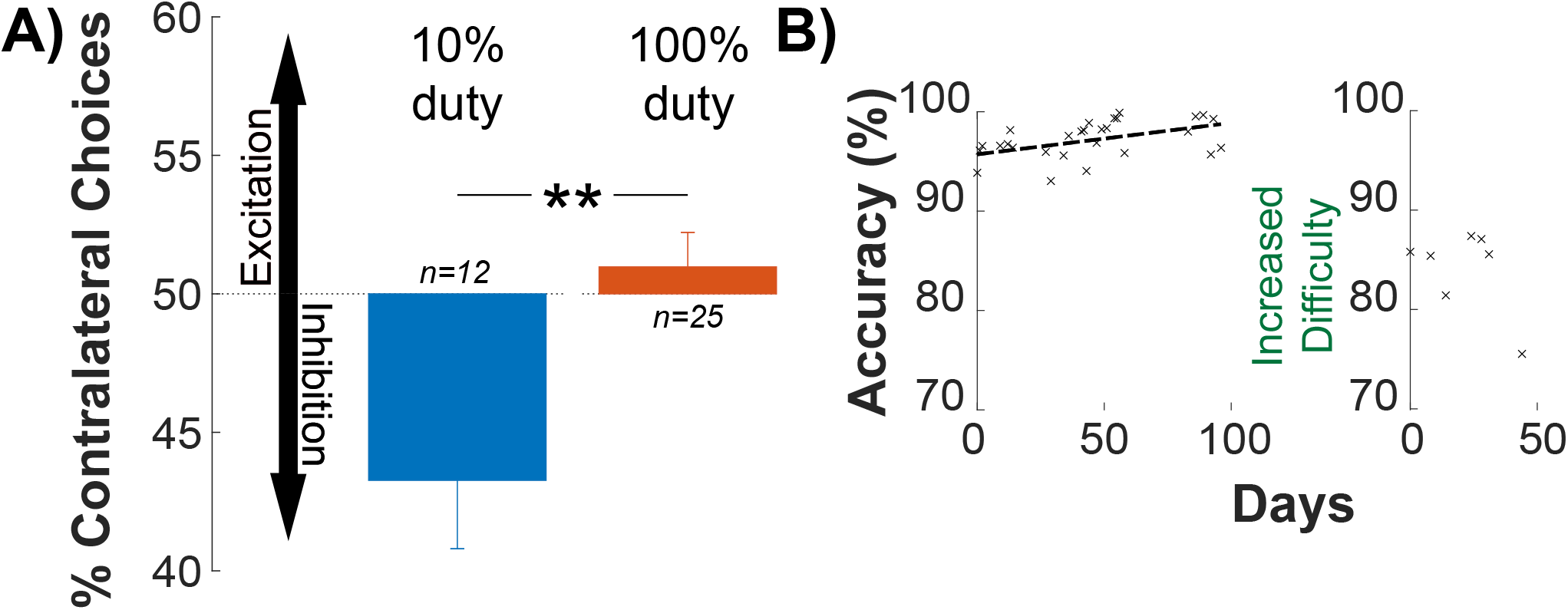
Inhibition of deep brain nuclei with immediate effects on choice behavior. **A)** Brief, low-intensity ultrasound (300 ms, 650 kHz, 1 MPa pressure at target) was delivered into the left or right LGN every 6-14 s. The side of the stimulation was randomized. The animal was deciding whether to look at a left or a right target. The brief, low-intensity ultrasonic pulses influenced the choice behavior. Specifically, variation in the duty cycle of the ultrasound significantly changed the proportion of contralateral choices. Stimuli pulsed at 10% duty significantly decreased the proportion of contralateral choices relative to a continuous stimuli. The decrease in contralateral choices is consistent with neuronal inhibition (vertical arrows). : *p <* 0.01. **B)** Repeated application of neuromodulatory ultrasound to deep brain circuits is safe. Significant damage to the LGN would cause a notable decline in the subject’s visual discrimination performance. We observed the opposite: the subject’s discrimination accuracy improved from day to day. This demonstrates that the repeated neuromodulation was safe at the functional level. The positive slope of the linear fit is significant before the increase in difficulty (*r* = 0.27, *p* = 0.004).

We found that stimuli pulsed at 10% duty cycle induced a significant (one-sided t-test, *t*(11) = −2.72; *p* = 0.02) ipsilateral bias, consistent with an inhibition or disruption of the neural activity in the target circuit. On the other hand, continuous stimuli induced did not have a significant effect on the animal’s bias during the task (one-sided t-test, *t*(24) = 0.78; *p* = 0.44). Notably, in this analysis, we only compared effects for trials in which there always was an ultrasound stimulus: either the left or the right LGN was sonicated. This controls for potential generic artifacts that can be associated with ultrasound. Moreover, we found a dissociation in the effect size by the stimulus duty cycle (two-sided t-test, *t*(35) = −3.1; *p* = 0.004). Intriguingly, the stimulus that delivered 10 times less energy into the target (10% duty) produced an effect while the more potent 100% duty stimulus did not significantly alter the animal’s behavior (Figure 2A). This corroborates the notion that duty cycle constitutes a critical variable in the neuromodulatory effects of ultrasound^29, 30^, and the growing consensus that the effects of low-intensity ultrasound are of mechanical, as opposed to thermal, nature^31–34^.

We delivered the ultrasound into the deep brain targets repeatedly, over 37 sessions. This enabled us to test the long-term safety of ultrasonic neuromodulation. Specifically, damage to the LGN would manifest as a decrease in the subject’s accuracy^35–38^. We found the opposite (Figure 2, bottom). The subject’s discrimination accuracy kept increasing over the course of the study (*p* = 0.004, line fit to the data in Figure 2B). The positive slope demonstrates the subject’s improvement in task performance over time. Near the end of the study we decreased the delays between the onset of the first and second target to make the task more difficult for the subject. While there is insufficient data after the increased difficulty to fit a line, there is no clear drop off in the subject’s performance. Thus, the subject’s behavior reveals no evidence of damage to either LGN.

Repeated delivery of ultrasound into the brain rests on reproducible positioning of the device and on quality coupling of the transducer face to the subject’s skin. To address these critical aspects of ultrasound delivery, we equipped the platform with imaging functionality that ensures reproducible positioning with respect to the skull and high-quality coupling (Figure 3). Specifically, the device has a receive capability that also implements pulse-echo imaging. The ultrasound images acquired during each session are compared with a standard taken during an MRI imaging session, which validated the targeting. In addition, the magnitude of the received echoes (Figure 3, bottom) provides information on the coupling quality. The resulting ultrasound image allows the operator to confirm repeatable and accurate placement of the array relative to the skull in each session but the image should not be interpreted as a visualization of skull thickness or brain anatomy. The measured variation in distance averaged across all 256 elements and 36 sessions (the coupling check failed to run in one session) was 0.19±1.0 mm. The variation in echo magnitude was -10±5.5%.

**Figure 3.**
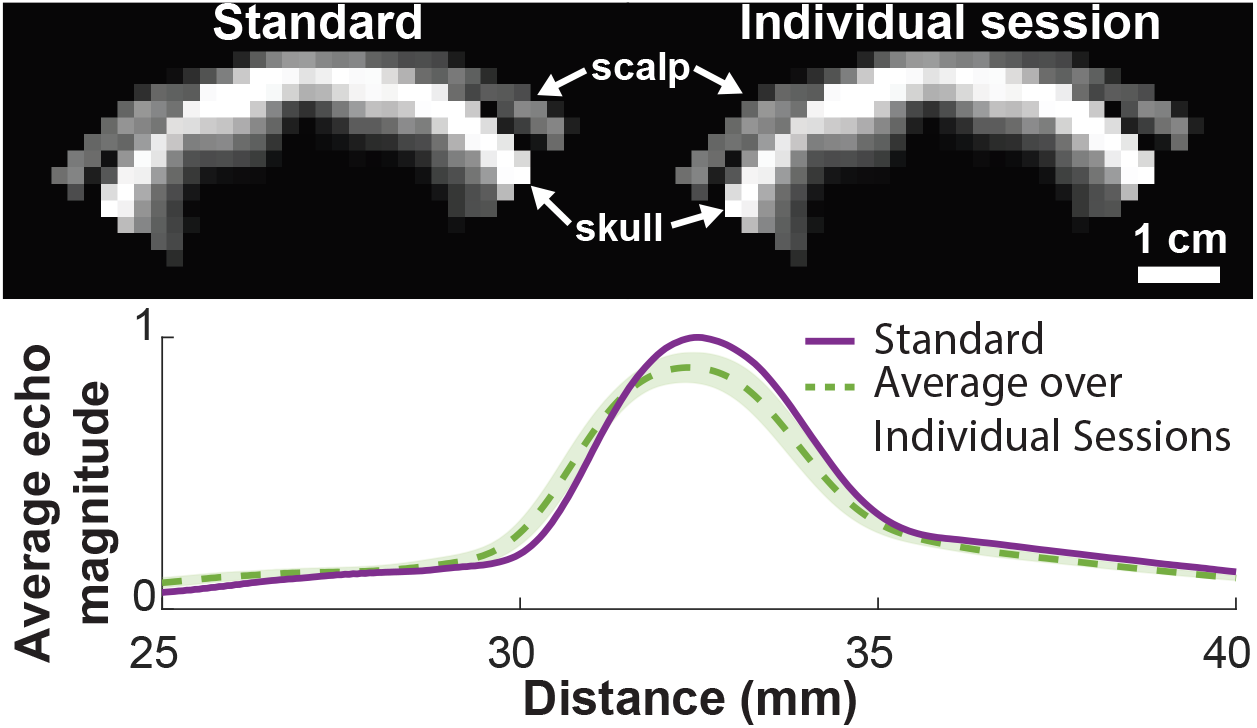
The system operated in imaging mode to ensure reproducible positioning and coupling. During each session, the platform is operated in imaging mode. **Top:** The imaging reliably detects the location of the scalp and the skull. **Bottom:** The timing of the echoes received from the skull provide information on the position of the device with respect to the skull; their magnitude evaluates the quality of ultrasound coupling to the subject’s head. Shaded region represents standard deviation across sessions, measurements acquired over 36 sessions.

Targets in the visual system, such as the LGN, will enable researchers to characterize the effects of ultrasound on neural tissue but future diagnostic and therapeutic protocols will require the ability to target systems involved in emotional regulation, such as the limbic system. Thus, we further validated our setup by demonstrating successful sonication of the LGN and amygdala in a second NHP subject (Figure 4).

**Figure 4.**
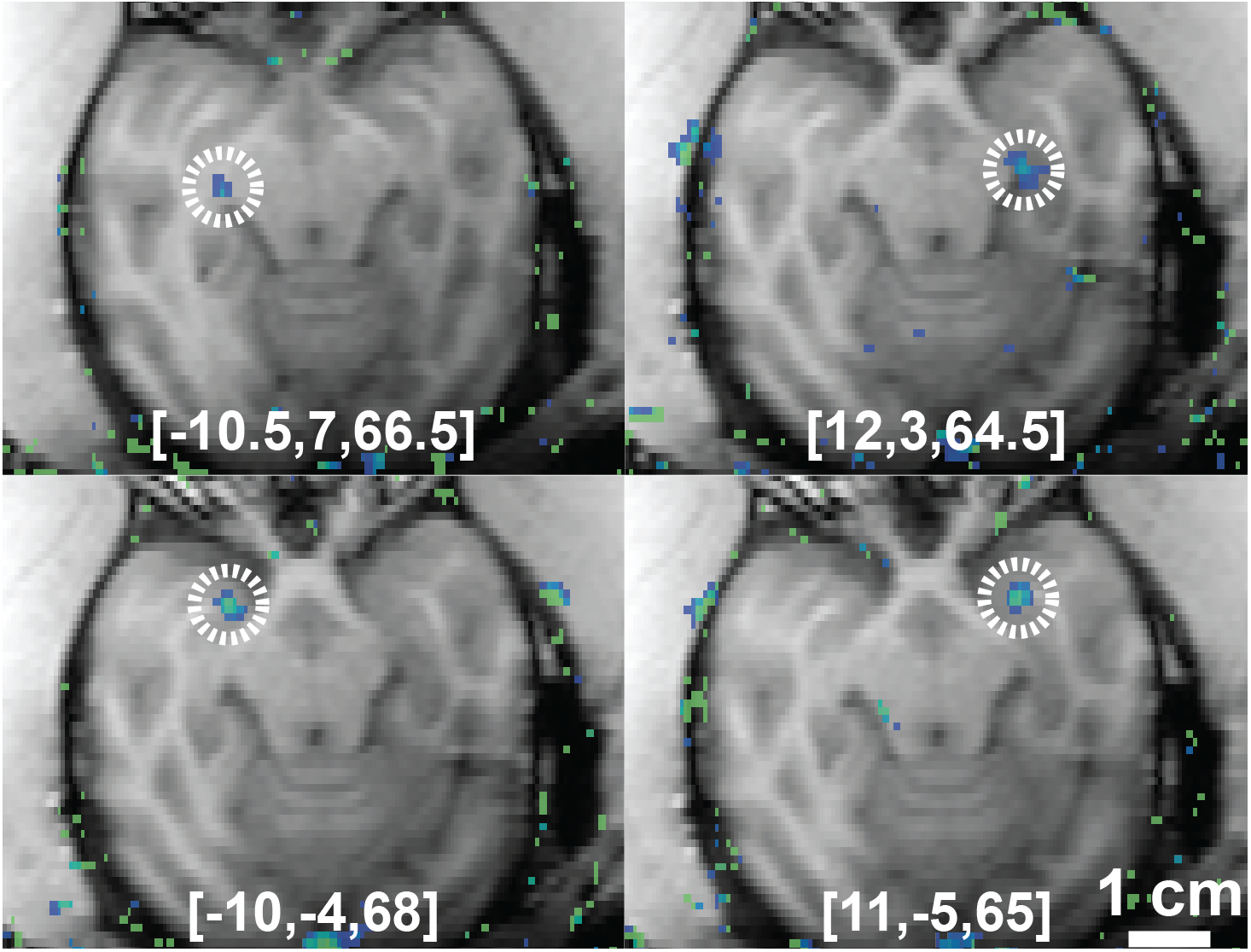
Flexible and Programmatic Deep Brain Targeting. Successful sonication of the left and right LGN and left and right amygdala is shown in a second NHP subject. The coordinates used to target each region are listed at the bottom of each image. These results demonstrate the capacity to reach a broad range of targets, including nuclei within the limbic system such as amygdala.

## Discussion

We developed and validated a practical and MRI-compatible system that delivers ultrasound through the intact skull and skin into one or more neural targets in awake subjects. Remus unleashes the full potential of ultrasound: personalized, targeted, and noninvasive interventions deep in the brain (Figure 5).

**Figure 5.**
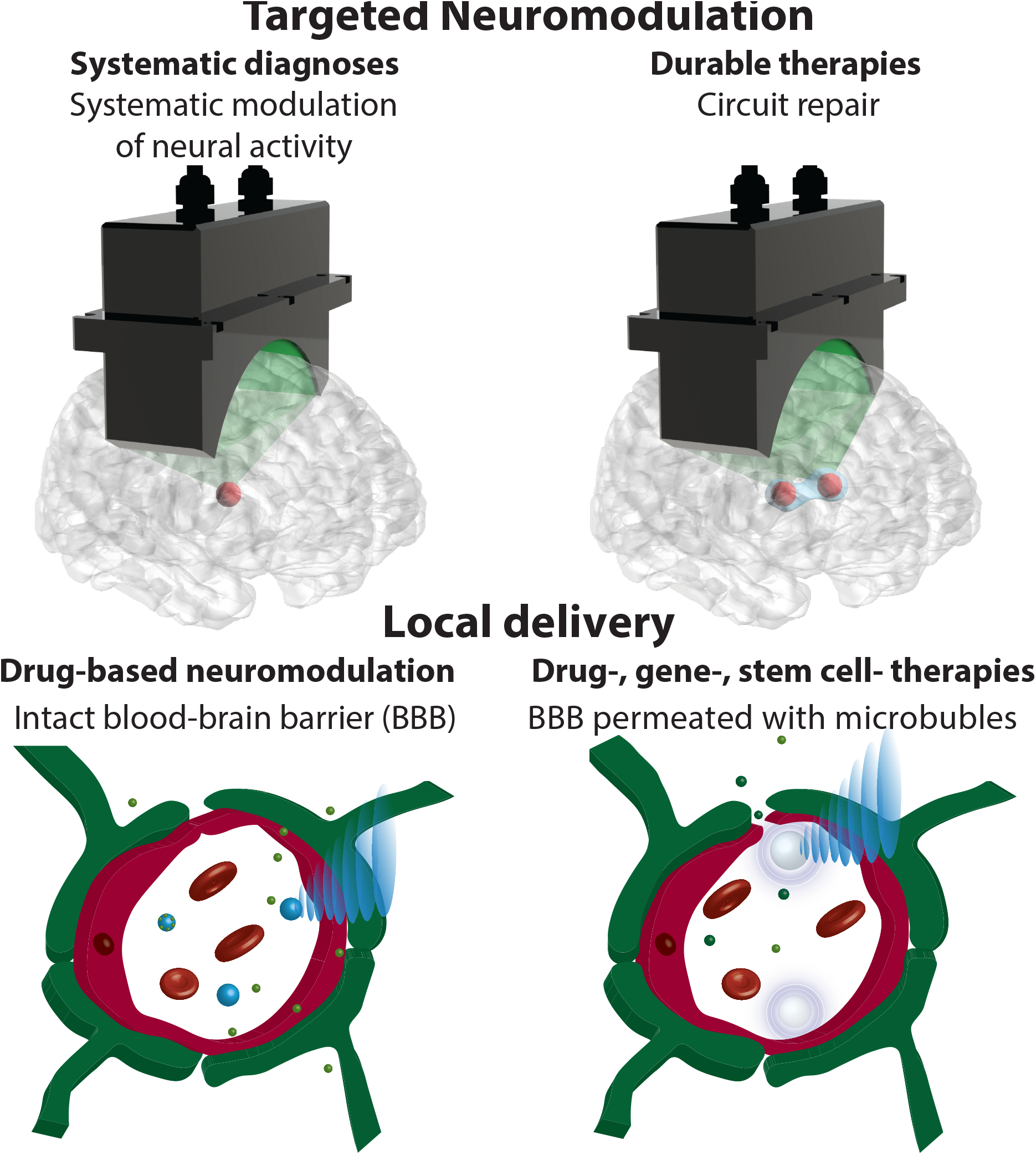
Applications of the platform for novel treatments of brain disorders. Brief, millisecond-second ultrasound stimuli modulate neural activity in a transient fashion. This effect, when coupled with successive targeting of individual circuit candidates, provides a tool to dissect circuit function and dysfunction (top left). Ultrasound delivered into a target for minutes induces neuroplastic changes in the target. This could be used for durable reset of the malfunctioning circuits (top right). To further increase the specificity of these effects, ultrasound can be combined with drug-carrying nanoparticles that release their cargo specifically at the ultrasound target (bottom left). If agents that do not naturally cross the blood-brain barrier (BBB) are to be delivered into a target, ultrasound can be combined with microbbubles that—upon sonication—transiently permeate the BBB (bottom right).

Remus provides this intersection of features as a portable device. On the other hand, emerging systems based on single element transducers are incapable of electronic targeting and have limited axial resolution. Surgical systems (ex: ExAblate Neuro, Insightec^26^) can deliver ultrasound into deep brain targets electronically, but the system was not designed for repeated, systematic applications. Moreover, in comparison with these systems, Remus provides imaging capabilities that validate the device’s position with respect to the head and test the quality of the ultrasound coupling.

The programmatic targeting enables clinicians to flexibly target multiple neural structures in a single session and the geometric design enables flexible mechanical placement of the array, resulting in a large treatable volume within the brain. This large treatment envelope, combined with the ease of targeting multiple nuclei in a single setting, provide the capacity to identify specific brain regions involved in an individual patient’s symptoms. For example, a physician can target multiple deep brain structures in a patient presenting with chronic pain. A significant change in the patient’s pain score would establish a causal relationship between the targeted region and the patient’s pain, thus enabling the physician to replace systemic treatment (such as opioids) with a more effective and targeted therapy.

The dissociation between neuromodulatory effects and duty cycle (Figure 2A) provide additional evidence of ultrasonic modulation of neural structures^31, 33^. In addition to these effects, ultrasound can elicit auditory or vestibular confounds^39, 40^. Remus’ fixed positioning and its capacity to programmatically deliver distinct ultrasound waveforms into distinct regions, with dissociated effects, enables scientists to control for such confounds.

### Future Applications

The manipulation of circuit function and dysfunction in the short term enables systematic and personalized diagnoses of the neural sources of brain disorders in the clinic. In the lab, transient manipulation of circuit function enables causal investigations of the function of specific deep brain regions in humans. Moreover, the system will enable researchers to deliver stimuli of durations necessary to induce circuit reset^15–17, 41^ into the malfunctioning regions.

Remus provides maximal pressure output over 5 MPa, well above the pressures needed to release drugs into a specified neural circuit during awake behavior. The awake setup enables investigators to quantify the effectiveness and duration of the neuromodulatory effects induced by specific neuromodulatory drugs, such as propofol^42^ or ketamine. In addition, the behavioral readout can be used to assess the safety of the release at the functional level (Figure 2B).

Analogous protocols can be used to assess the effectiveness and safety of cargo delivery across the blood-brain barrier (BBB)^21^. To maximize therapeutic effects, agents, such as chemotherapy, need to be delivered across the BBB repeatedly over multiple sessions. This may raise safety concerns^43^. Remus can be used for systematic investigation of the effects of repeated ultrasound-based BBB opening.

Together, Remus affords multi-focal ultrasound-based brain intervention at high spatiotemporal resolution. The system’s flexible targeting and its validation of positioning and coupling open new possibilities for the large number of patients currently without options.

## Methods

### Animals

Two male macaque monkeys (*macaca mulatta*) participated in this study. One subject (age 6 years, weight 8 kg) participated in both MR based targeting validation and behavioral studies. A second subject (age 6 years, weight 12 kg) participated only in MR based target validation. The procedures were approved by the Institutional Animal Care and Use Committee of the University of Utah.

### System Overview

An overview of the system is given in Figure 6. Custom targeting software interfaces with the MR scanner for target validation and with the ultrasound driving system to deliver ultrasound to the desired location. Thus, Remus delivers ultrasound into specific deep brain targets, enabling real-time modulation of awake behavior. Items in the green boxes are provided as part of this study at onetarget.us/software.

**Figure 6.**
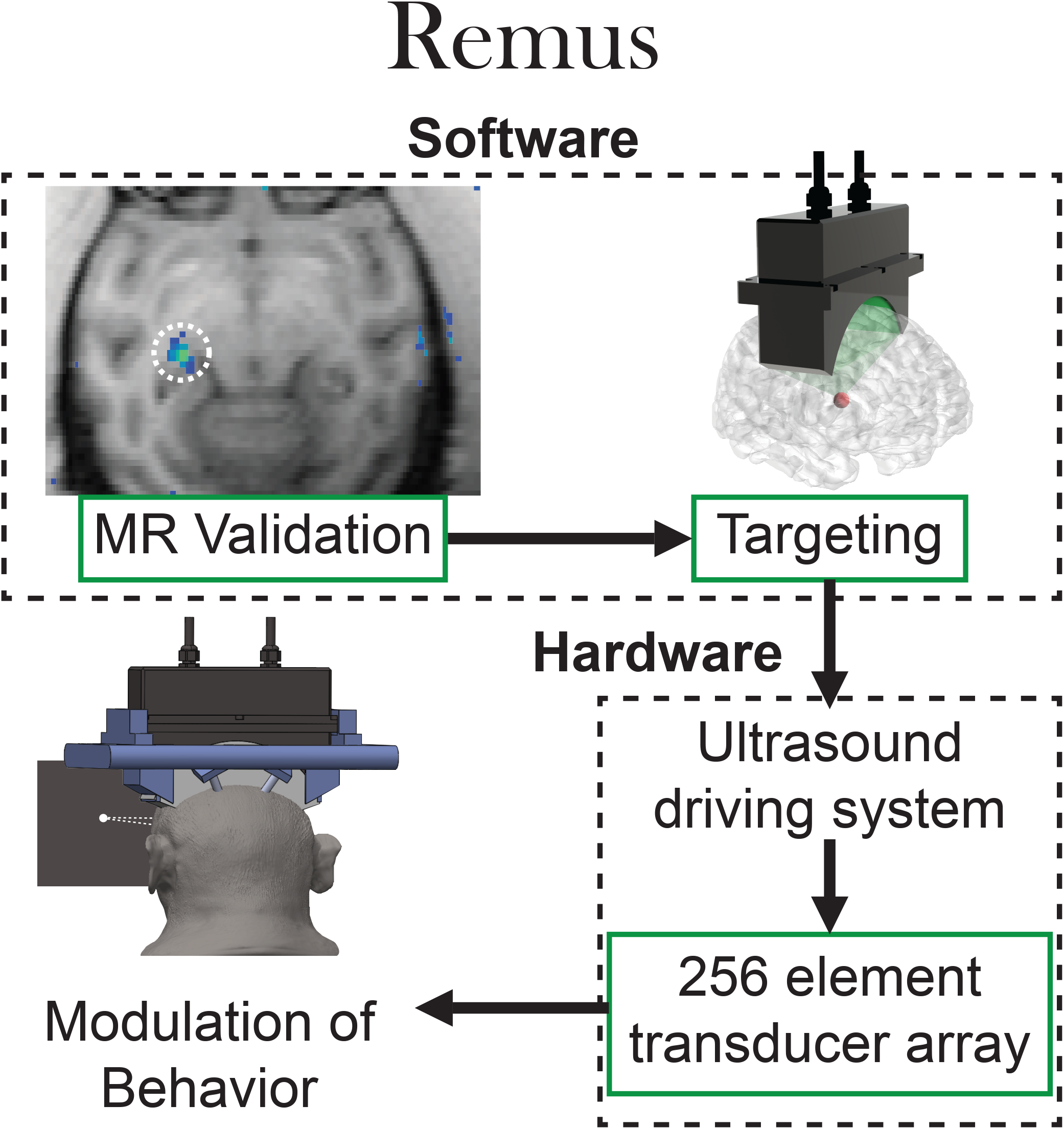
Overview of Remus. Custom targeting software communicates with the MR scanner to validate targeting of the desired neural-anatomy. The custom MR software overlays the ultrasound induced temperature rise on anatomical images, allowing the user to adjust targeting for optimal coverage of the desired region. The targeting software then controls the Verasonics system which drives the 256 element array. The elements in green boxes can be found at (onetarget.us/software).

#### MR Validation

A series of custom MATLAB functions allows the user to register the array to an MR image, select an acoustic focus within the image, and deliver a sonication to the desired anatomical location. The resulting thermometry images are overlaid on the anatomical image to verify accurate placement of the acoustic focus.

#### Targeting

Custom targeting software interfaces with the ultrasound driving system (Verasonics, Kirkland Washington) to measure the distance between the array and the skull, measure the quality of the acoustic coupling, and to provide the individual element phases necessary to place the acoustic focus at the desired location. Phases are determined using a time of flight algorithm^44^.The targeting software accepts user inputs through a serial connection to enable a user to specify target locations and to change those locations during individual sessions.

#### Cost

The cost of Remus is dictated by the costs of the driving system and the transducer array. Bulk manufacturing of the complete system is currently estimated to be around $50,000.

### NHP Platform Positioning and Head Fixation

For the NHP experiments shown here, we developed an apparatus that ensures reproducible positioning of the platform with respect to the head from session to session, and at the same time allows us to head-fix NHPs and engage them in behavioral tasks. To achieve that, we developed a custom head frame that is attached to the skull via four titanium pins (Gray Matter Research, Bozeman, MT). Each pin is attached to the skull using three screws. The frame for the NHP experiments was designed in 3D cad software and produced with a 3D printer. A 3D drawing of the frame is available at onetarget.us/software. The frame is attached to a primate chair using two steel bars, mounted into the left and right side of the frame. Coupling of the transducer to the skull is achieved using a 6% polyvinyl alcohol (PVA) gel^45^. The subject’s hair is shaved prior to each session. Shaving may not be necessary in the ultimate human applications; the platform could detect the quality of the coupling and aberration due to the hair, and adjust the emitted ultrasound amplitude accordingly.

In each session, the system measures the accuracy of the device placement with respect to the head and the quality of the acoustic coupling using ultrasound imaging. Specifically, the system measures the distance between the transducer and the skull at six distinct locations. A pulse-echo measurement, performed on a 3×3 grid of elements, provides an estimate of the location of the skull relative to the transducer. The distance is measured by detecting the front edge of the reflected ultrasound energy and assuming a speed of sound of 1500 m/s in the coupling gel. The average difference in position relative to the position measured during MR thermometry was 0.47 ± 0.55 mm. The total power returned to each element (Figure 3, bottom) was 9 ± 8% lower across sessions compared to the power measured during the MR thermometry session.

### MR Imaging

The system is fully MRI-compatible. Accurate targeting of specific deep brain regions can therefore be validated using MRI, such as MR thermometry or MR ARFI. We used MR thermometry. In this approach, a continuous, 5-second sonication is sufficient to increase temperature at the focal spot by about 2^*°*^C, enabling visualization of the focus without inducing long-term changes in the neural tissue.

All scans were performed using a 3T MRI scanner (Trio, Siemens Medical Solutions, Erlangen, Germany) with a 4-channel flex coil wrapped underneath the animal’s head. High resolution 3D T1-weighted images were used for anatomical imaging and target identification; Magnetization Prepared Rapid Acquisition Gradient Echo (MPRAGE), TR=1900 ms, TE=3.39 ms, TI-900 ms, BW=180 Hz/pixel, flip angle 9^*°*^, FOV = 192×192×120 mm, resolution 1×1×1 mm, acquisition time 5:12 min. MR thermometry was performed with a 3D gradient recalled segmented echo planar imaging pulse sequence; TR=24 ms, TE=11 ms, BW=592 Hz/pixel, flip angle 12^*°*^, echo train length = 5 with monopolar readout, FOV = 144×117×36 mm, resolution 1.5×1.5×3.0 mm, acquisition time 4.6 s per dynamic image.

Registration of ultrasound and MR space was performed using three fiducial markers. After registration, the average euclidean distance between the estimated and the actual location of of the three fiducials (as measured by the 3-dimensional center of mass) was 2.5 mm. The transducer caused distortion along the subject’s inferior/superior axis resulting in approximately 4 mm of error in the transducers registration in that dimension. The registration of the transducer to the subject’s anatomy guided initial steering of the acoustic beam. Once the first thermometry signal was acquired, adjustments to the focal location were based primarily on the location of the thermometry signal, thus correcting any errors introduced by poor transducer localization.

It is well established that preferential absorption of ultrasound in the skull can lead to undesirable off-target heating. Indeed, clinical ablation trials rely on both a large transducer aperture and constant cooling of the scalp to mitigate this risk^26^. With a smaller transducer aperture and in the absence of active cooling investigators must assume that the temperature rise in the skull could be higher than at focus. Care should thus be taken to monitor the health of the skull and scalp between the transducer and the target when MR thermometry is used to validate acoustic targeting.

At the short durations and low frequencies used for neuromodulation in this study, heating of the skull was not a concern. The subject, who was awake, showed no signs of discomfort in response to the ultrasound.

### Acoustic Intensity at Target

The MRI thermometry additionally allowed us to compute the ultrasound pressure delivered in the target. Assuming negligible conduction and perfusion and a continuous sonication, the acoustic intensity, *I*, is related to the temperature rise, Δ*T*, by^46^

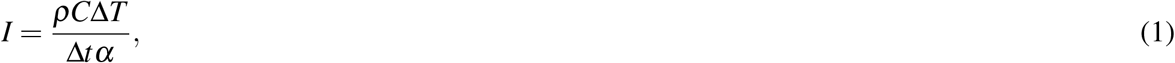

where *ρ, C*, and *α* are the density, specific heat capacity, and acoustic absorption of the tissue and Δ*t* is the time in which the temperature increase occurs. To minimize the effects of conduction and perfusion Δ*T* is chosen to be the maximum temperature measured in the seventh dynamic (the first dynamic in which the ultrasound is on). The center of k-space is acquired 2.3 seconds into each acquisition. Thus, Δ*t* was set to 2.3 s. We assumed a density of 1046 *kg/m*^3^, a specific heat capacity of 3630 J/K, and an acoustic absorption of 3.9 Np/m.

The MR thermometry can also be used to estimate the maximum temperature rise during each neuromodulatory sonication. The temperature rise during the first 2.3 s of each thermometry pulse was 1.4 and 1.8 C in the left and right LGN respectively. The 10% duty cycle sonication delivered 1.3% of the energy delivered in the first 2.3 seconds of the thermometry sonication. Thus, we can estimate a temperature rise of 0.014 and 0.018 C in the left and right LGN. A continuous wave sonication delivers 10 times the energy, thus, the expected temperature rise for these sonications is 0.14 and 0.18 C.

### Stimulation Parameters

Our custom, 256 element transducer (Doppler Electronic Technologies, Guangzhou, China) can produce focal pressures greater than 3 MPa at its 650 kHz center frequency and a usable freuquency range of 480-850 kHz. The transducer geometry is semicircular, with a radius of curvature of 65 mm (see Figure 1). The elements are 4 × 4 mm square and the distance between elements is 4.2 mm. All stimuli used 650 kHz. The pulsed stimulus (10% duty) used a 500 Hz pulse repetition frequency. Both pulses used a square envelope. The peak focal pressure (estimated from the thermometry data) was 1.1 and 0.98 MPa in the right and left LGN respectively. The half power beam width measured by a hydrophone in a water tank for was 1, 3.75, and 3.75 mm in the left/right, anterior/posterior, and superior/inferior dimensions, respectively. The half power beam width measured by thermometry was 0.75 and 2.25 mm in the left/right dimension and 3.75 and 4.5 mm in the front/back dimension for the left and right LGN respectively. Using the same input voltage, the free field pressure–measured using a hydrophone–is 2.4 MPa at the location of the left and right LGN. Thus, the MR thermometry measurements suggest that about 30% of the pressure reaches the target through the individual layers. It is worth noting, nonetheless, that this estimate likely underestimates the actual pressure at target. In particular, the temperature measured by the MR is averaged spatially (across a voxel) and temporally (across the acquisition time); the actual peak temperature is thus higher than the measured average. In addition, a portion of the energy is distributed to the thermal conduction and vascular convection.

### Task

We trained one NHP to perform a visual choice task. This task was used in many previous studies (e.g.,^10, 47^). Briefly, the subject was seated in a comfortable primate chair and was presented with visual stimuli shown on a monitor. The subject’s eye movements were tracked and recorded using an infrared eye tracker (Eyelink, SR Research, Ottowa, Canada). In the task, the subject first fixates a central target. Following its offset, one target appears on the left and one target on the right side of the screen. There is a brief, controllable delay between the onset of the two targets, which can range from -40 to 40 ms. We varied the location of the targets within 2.5 visual degrees to circumvent adaptation. The sonication of the left and right LGN was randomly interspersed with trials in which no ultrasound was delivered. In an ultrasound trial, a 300 ms stimulus was delivered 150 ms before the fixation point offset. The subject receives a liquid reward if he looks at the target that appeared first within 2 s. In the key condition in which both targets appear at the same time and during which we quantify the effect of ultrasound, the subject is rewarded for either choice. The subject generally worked daily throughout the work week.

### Software

We designed custom software to integrate ultrasound sonication with both the MR thermometry and the behavioral task. The software can register the transducer to an MR coordinate system, output the targeted focus in both transducer and MR coordinates, and overlay the temperature change (as measured by MR thermometry) on the anatomy image. To protect the subject’s scalp, skull, and neural tissue we used very small temperature increases to visualize the focal spot. Thus, to improve the visualization, we included a noise reduction algorithm in our visualization software. The algorithm assumes an exponential increase in temperature during the sonication and an exponential decay in temperature after the ultrasound turns off. Voxels in which the temperature does not follow such a trend are ignored.

Similarly, we developed software that integrates a behavioral task with flexible sonication of the targeted neural circuit. The ultrasound controller accepts target coordinates and intensities from the server running the behavioral task. Synchronization of the sonication with presentation of the visual stimulus is achieved with a TTL pulse. All of the ultrasound software was developed within the Verasonics framework and is available at onetarget.us/software.

## Acknowledgments

This development of the platform and its application in NHPs was supported by the National Institute of Neurological Disorders and Stroke, Grants 5R00NS100986 and F32MH123019. We thank Tyler Davis and Eric Burdett for facilitating this work.

